# Parameter Estimation in Brain Dynamics Models from Resting-State fMRI Data using Physics-Informed Neural Networks

**DOI:** 10.1101/2024.02.27.582428

**Authors:** Roberto C. Sotero, Jose M. Sanchez-Bornot, Iman Shaharabi-Farahani

## Abstract

Conventional modeling of the Blood-Oxygen-Level-Dependent (BOLD) signal in resting-state functional Magnetic Resonance Imaging (rsfMRI) struggle with parameter estimation due to the complexity of brain dynamics. This study introduces a novel brain dynamics model (BDM) that directly captures BOLD signal variations through differential equations. Unlike dynamic causal models or neural mass models, we integrate hemodynamic responses into the signal dynamics, considering both direct and network-mediated neuronal activity effects. We utilize Physics-Informed Neural Networks (PINNs) to estimate the parameters of this BDM, leveraging their ability to embed physical laws into the learning process. This approach simplifies computational demands and increases robustness against data noise, providing a comprehensive tool for analyzing rsfMRI data. Leveraging the functional connectivity matrices scaled by the estimated parameters, we apply a state-of-the-art community detection method to elucidate the network structure. Our analysis reveals significant differences in the participation coefficients of specific brain regions when comparing neurotypical individuals to those with Autism Spectrum Disorder (ASD), with distinct patterns observed between male and female cohorts. These differences are consistent with regions implicated in previous studies, reinforcing the role of these areas in ASD. By integrating PINNs with advanced network analysis, we demonstrate a robust approach for dissecting the complex neural signatures of ASD, providing a promising direction for future research in neuroimaging and the broader field of computational neuroscience.

## I. INTRODUCTION

The study of brain dynamics through the Blood-Oxygen-Level-Dependent (BOLD) signal measured by functional Magnetic Resonance Imaging (fMRI) has been crucial in advancing our understanding of cognitive functions and neurological disorders [1], [2]. Foundational modeling frameworks such as the Balloon model [3], [4], Dynamic Causal Modeling (DCM) [5], [6], the metabolic/hemodynamic model (MHM) [7], and neural mass modeling [8], [9], have furthered our understanding of signal linked to neuronal activity. However, these models often require extensive a priori knowledge and struggle with parameter estimation challenges arising from the noise and complexity inherent in fMRI data [10]. When modeling extensive brain networks, the high dimensionality of such models leads to considerable computational challenges in fitting them to empirical data [5], [11]. Additionally, frameworks like DCM and MHM were designed for task-based fMRI data because of the complexities involved in fitting brain dynamics models to resting-state fMRI (rsfMRI) data. Recently, the DCM framework was extended to accommodate rsfMRI data, employing spectral methods that focus on frequency domain representations and retain only linear terms, potentially neglecting the rich non-linear interactions present in the brain’s intrinsic activity [6], [12].

The development of Physics-Informed Neural Networks (PINNs) [13] has provided a robust solution for parameter estimation, integrating physical laws into the training of deep learning models. PINNs are particularly useful in fields where conventional data-driven approaches may fall short due to limited or noisy data. By incorporating biological laws and constraints within the PINN framework, they can effectively solve a range of differential equations, including partial differential equations (PDEs), integral-differential equations, and stochastic PDEs [14], making them highly suitable for studying brain dynamics models (BDMs).

In the context of this paper, PINNs are applied to present a novel method that utilizes a scaled functional connectivity matrix to enhance the parameter estimation process for fMRI data. This methodological advancement simplifies computational analysis and effectively captures the unique dynamics of each brain region. By leveraging the PINN framework, the proposed approach can streamline complex modeling tasks, potentially leading to more accurate and efficient analysis of neural data. Specifically, we introduce a nonlinear BDM of the BOLD signal that scales interactions with other brain areas as provided by the functional connectivity matrix. Using this model, we construct a PINN and employ it to concurrently estimate the parameters of BDM while training the network, offering a new tool for probing the mechanisms underlying brain activity patterns in neurotypical and autism spectrum disorder (ASD) subjects.

## II. METHODS

### A. rs-fMRI data preprocessing and preparation

In this study, we analyzed rsfMRI data from the Autism Brain Imaging Data Exchange (ABIDE) initiative registered in 17 international sites [15]. The dataset comprises 1112 subjects, of which 539 have ASD (474 males and 65 females), and 573 are neurotypical (474 males and 99 females). For the ASD group, the average age of males is 17.14±8.43, and the average age of females is 16.09±7.82. For the neurotypical group, the average age of males is 17.43±7.90, and the average age of females is 15.44±6.57. More detailed demographics information is presented in [16]. To ensure consistency and avoid the variability introduced through different preprocessing pipelines, we opted for datasets preprocessed with the Configurable Pipeline for the Analysis of Connectomes (CPAC), as provided by ABIDE. The details for this preprocessing procedure are provided in the literature [17].

We acknowledge the ongoing debates regarding the optimal preprocessing techniques for rsfMRI data, particularly the use of bandpass filtering and global signal regression (GSR). Since opinions on these methods vary, and outcomes can be contingent upon the chosen features and neural network architectures [18], [19], we implemented two commonly accepted preprocessing steps: bandpass filtering within the range of 0.01 to 0.1 Hz and GSR. To address the variability in BOLD-rsfMRI data from different ABIDE collection sites [20], we implemented a standardization process. To begin, we resampled the time series from each participant’s brain area to a resolution of 2 seconds. Subsequently, these time series were segmented into intervals of 3 minutes. Finally, we standardized each BOLD time series by applying Z-score normalization.

To construct functional connectivity matrices for each subject, Pearson correlation coefficients were computed between time series from the 116 brain regions defined by the Automated Anatomical Labeling (AAL) atlas [21]. In assessing the significance of these connections, we employed the False Discovery Rate (FDR) [22] to correct for multiple comparisons, setting connections with FDR-adjusted p-values above 0.05 to zero, thus excluding them as non-significant.

### B. rsfMRI-based Brain Dynamics Model

Our goal is to construct a model for the BOLD-rsfMRI time series data of each brain region, using a differential equation framework that accounts for inter-regional interactions. Within this model, the time series for the i-th brain area, denoted as *y*_*i*_ (*t*), is defined as follows:

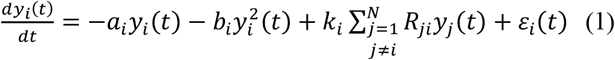

The term = ™*a*_*i*_*y*_*i*_ (*t*) with *a*_*i*_> *0* represents linear negative feedback, ensuring the signal damping which is a common homeostatic feature in physiological systems [23]. The nonlinear term 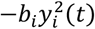 counterbalances the grow of *y*_*i*_ (*t*) thereby introducing a factor that captures the stabilizing effects observed in neural responses as activities reach higher levels [24]. The summation term 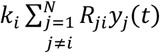 embodies the collective influence of all other brain regions on the i-th region, weighted by the functional connectivity matrix *R*_*ji*_, which is derived from empirical data. The coefficient *k*_*i*_scales this influence and can be interpreted as the degree of receptiveness of region *i* to inputs from other areas. This model is a simplification yet captures essential features of brain dynamics, like signal propagation and feedback mechanisms. Finally, *ε* _*i*_ (*t*) are random perturbation that are considered as uncontrolled and independent for each area.

### C. The Physics-Informed Neural Network approach

PINNs represent a groundbreaking intersection of machine learning and physical sciences. PINNs integrate known physical laws into the structure of neural networks, allowing them to predict outcomes that adhere to these laws, thus offering a powerful tool for solving complex differential equations that are ubiquitous in modeling physical systems [13]. This fusion of physics and deep learning has profound implications across various fields, including biology [25], where the governing laws are expressed through differential equations that describe biological processes from molecular interactions to the dynamics of ecosystems.

The use of PINNs can be extended to the field of neuroscience, particularly in the modeling of brain activity as captured by fMRI data. Here, PINNs offer a novel approach for estimating the parameters of brain dynamics models. The key advantage of PINNs in this context is their ability to constrain the solution space to feasible dynamics that are consistent with both the observed data and the underlying biological processes. This is particularly important in neuroimaging, where the data is high-dimensional and noisy, and the systems of interest are highly nonlinear and complex.

In this work, we harness the capabilities of PINNs to estimate the model parameters from rsfMRI data, using the framework provided by the DeepXDE Phython library [26]. Specifically, we estimate the parameters of the BDM for each participant and their respective brain regions. As depicted in Fig. 1 through a flowchart, the process initiates with the neural network receiving time (t) as input and subsequently predicting the BOLD signal, *y*_*i*_ (*t*), for each specific brain area. The network’s performance is gauged by the Mean Square Error (MSE) loss function, which is an amalgamation of two distinct loss components: Data Loss and Physics Loss. Data Loss measures the deviation between the predicted BOLD signals and the actual recorded signals, while Physics Loss ensures the predicted signals conform to the dynamics stipulated by the BDM’s differential equation. The network strives to estimate the parameters within this differential equation to simultaneously fit the observed data and abide by the physical laws. The differential equation under consideration includes both linear and nonlinear terms, scaled by parameters *a*_*i*_ and *b*_*i*_, as well as a term signifying the cumulative effect of other brain areas, scaled by parameter *k*_*i*_ and the functional connectivity *R*_*i*_. This integration of physical laws into the learning process yields a model that is both data-centric and physically coherent, offering a more comprehensive and precise depiction of brain functionality. To simplify our model, we posit *ε* _*i*_ (*t*) as null, suggesting that signal variations stem solely from the inherent oscillations in the BOLD time series of the targeted area and its networked regions.

**Fig. 1.**
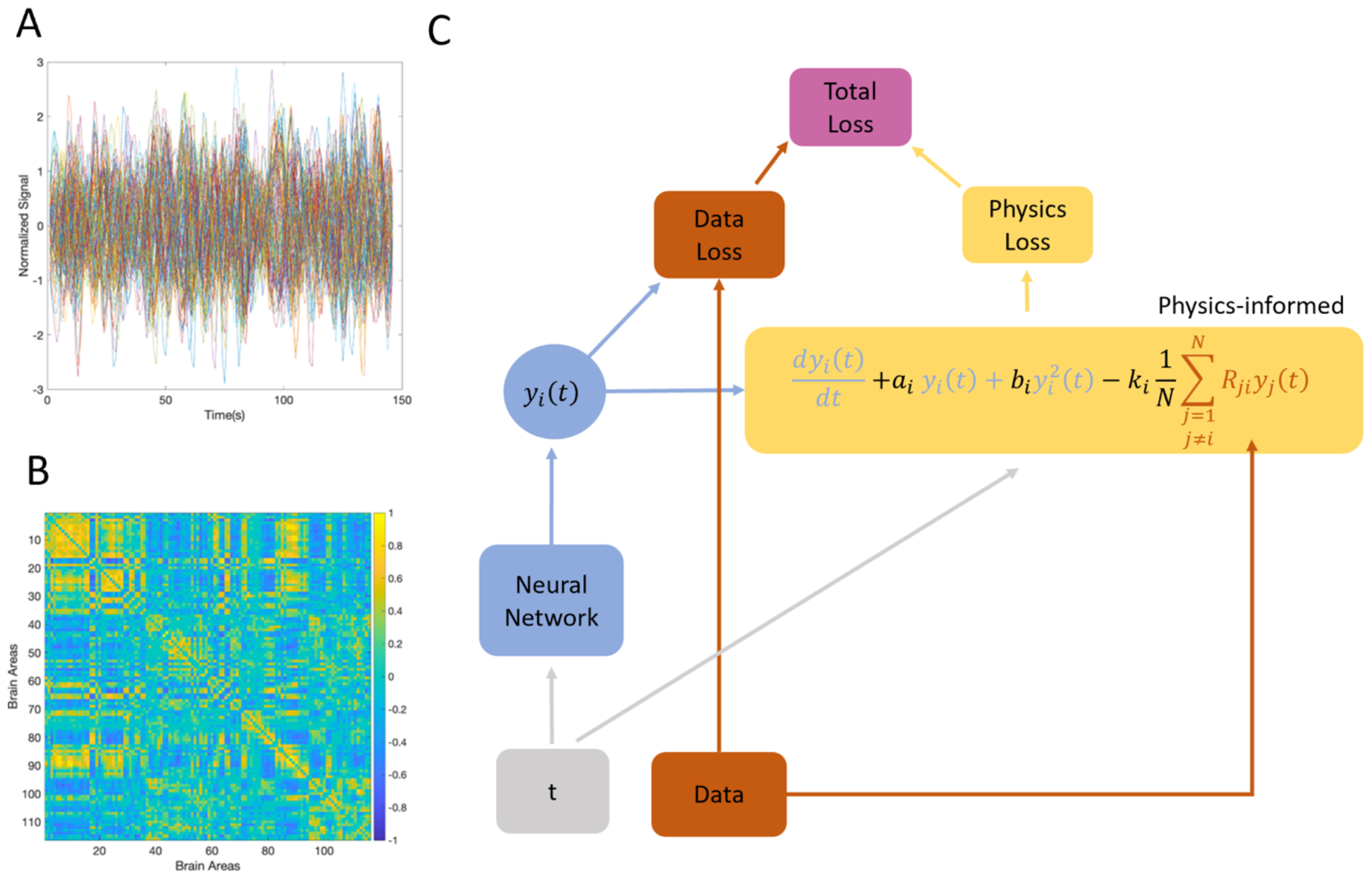
Schematic of the PINN for parameter estimation of a BDM using rsfMRI data. A) The normalized BOLD signals for the 116 brain regions considered for one subject. B) The Functional Connectivity Matrix *R*_*ji*_ for the same subject as in A, constructed with the Pearson correlation coefficients between the signal time courses of different brain regions. C) The model’s architecture is summarized in a flow diagram, beginning with the input time (t) to the Feedforward Neural Network that predicts the BOLD signal *y*_*i*_ (*t*) for each brain area. The Total Loss function is composed of Data Loss, which quantifies the discrepancy between the predicted and actual fMRI signals, and Physics Loss, which ensures the predicted signals adhere to the dynamics defined by the differential equation. The equation incorporates linear and nonlinear terms of the BOLD signal, scaled by parameters *a*_*i*_ and *b*_*i*_, and a term representing the average influence of other brain regions, scaled by the parameter *k*_*i*_ and the functional connectivity *R*_*ij*_, thus integrating the physical laws governing brain dynamics into the learning process.

The neural network in Fig. 1 is a fully connected feedforward neural network, structured with an input layer (time), three hidden layers of 100 neurons each, and an output layer that consists of a single neuron. This architecture was implemented using DeepXDE’s dde.nn.FNN function. The hyperbolic tangent (tanh) function served as the activation function, providing the necessary non-linearity for capturing complex brain dynamics. Additionally, we employed the Glorot uniform initializer for weight initialization, which is known to facilitate faster and more effective training convergence in deep neural networks.

In terms of training methodology, the network parameters are initially set using random values, ensuring a degree of variability in the starting conditions of the learning process. The training is conducted using the Adam optimizer [27], a popular choice for its adaptive learning rate capabilities, with a learning rate set at 0.001. This relatively low learning rate helps in a gradual and more stable convergence. The network is subjected to a substantial training regimen, running for 40,000 iterations. This extensive training is crucial for the network to adequately learn the complex dynamics of the rsfMRI data.

## III. RESULTS

Results in Fig. 2 show significant differences in the estimated parameters for certain brain regions when comparing individuals with autism to neurotypical controls. These differences are captured in the parameters *a*_*i*_, *b*_*i*_, and *k*_*i*_ which are integral to the modeling of brain dynamics using resting-state fMRI data. For the male cohort (row A), the bar graphs reveal a varied pattern of mean differences across multiple brain areas. The statistical contrast of neurotypical minus autism for the nonnegative parameter *a*_*i*_, which reflects the linear rate of change of the BOLD signal, shows both significant positive and negative differences across brain regions. This suggests that in some areas, the linear dynamic is more pronounced in the neurotypical group, whereas in others, it is more pronounced in the autism group. The parameter *b*_*i*_, indicative of non-linear dynamics, exhibits positive mean differences in regions such as the right Parietal Inf and left Putamen, suggesting a stronger non-linear response to fluctuations in BOLD signal in the neurotypical group within these areas. The scaling factor *k*_*i*_, which modulates the influence of connectivity, generally shows a higher mean in the neurotypical group, except for one region, indicating a potential difference in how brain regions interact within the network.

**Fig. 2.**
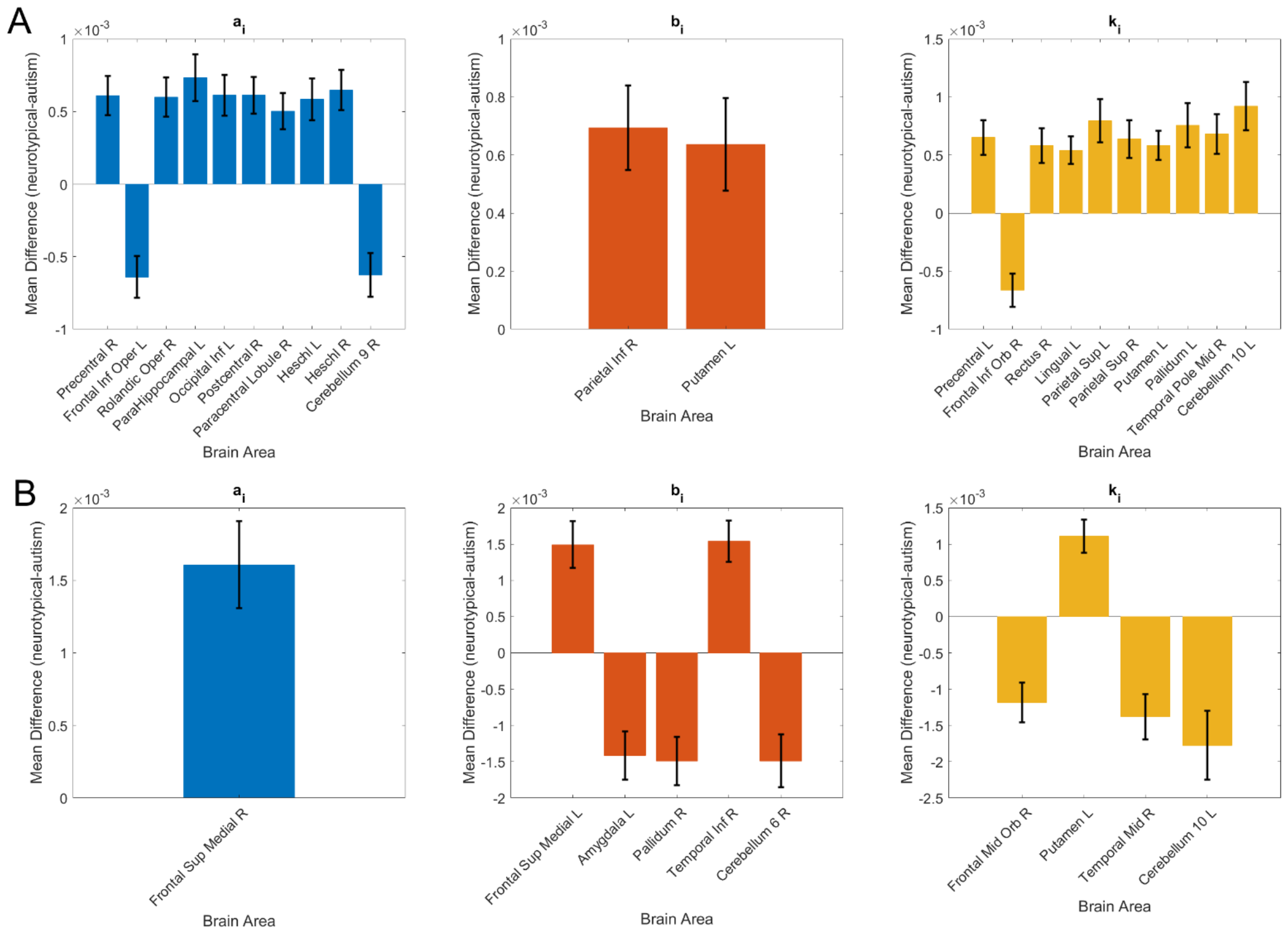
Gender-specific parameter estimation from rsfMRI data using PINNs. A) Male cohort: The bar graphs illustrate the significant mean differences in the estimated parameters *a*_*i*_, *b*_*i*_ and *k*_*i*_ across brain regions for the male group, comparing individuals with autism to neurotypical controls. Error bars indicate the standard error of these estimates, underscoring the variability within each group. B) Female cohort.

In the female cohort (row B), we only found one area with significant differences for the linear parameter *a*_*i*_ (as opposed to 11 areas for the male cohort) and 5 areas with significant differences for the nonlinear parameter *b*_*i*_ (as opposed to only 2 areas for the male cohort). This suggests a distinct linear and nonlinear dynamic in male and female cohorts. However, it should be noted that males and females present a different prevalence of ASD which is reflected in the larger number of males subjects than female subjects in the ABIDE dataset. The specific brain areas found to exhibit significant differences are notable. Previous studies have identified several of them as regions of interest in the context of autism. For example, the frontal lobe, and specifically the medial frontal cortex, has been implicated in autism, often associated with executive function and social cognition deficits commonly observed in individuals with the condition [28], [29]. The putamen, part of the basal ganglia, plays a role in motor planning and learning and has been shown to have altered activity in individuals with autism [30], potentially related to repetitive behaviors and motor anomalies.

Our analysis extended beyond evaluating the estimated parameters to examining the networks shaped by scaling the functional connectivity matrices with the estimated parameters *k*_*i*_. To study these networks we applied a recently proposed community extraction methodology optimized for correlation-based networks [31]. This methodology combines Hebbian learning with random walk-based exploration of the correlation network [32] and with the Louvain method for detecting communities within signed networks which is implemented in the Brain Connectivity Toolbox (BCT) [33]. The parameters were defined as follows: the length of the time series generated by the random walkers was set to *T* = 1000, the number of random walkers exploring the network was *K* = 10^−^, and the learning rate for the Hebbian learning rule applied during the random walk was *α* = *0*.5. To ensure robustness in our community detection, we replicated this process 40 times yielding an ensemble of 40 community partitions for each subject. A consensus partition was then achieved using the algorithm developed by Lancichinetti and Fortunato [34], which is implemented in the BCT. We utilized this consensus partition to calculate the participation coefficients from positive (Ppos) and negative (Pneg) weights. The participation coefficient describes the distribution of a node’s connections across communities [35] and ranges from 0 to 1. A coefficient approaching 1 suggests that a node’s connections are evenly spread among different communities, indicating integrative nodes. Conversely, a coefficient near zero signifies that most connections are confined to the node’s own community, highlighting provincial nodes. These measures are instrumental in understanding the role and affiliation of individual nodes within the broader network structure.

Fig. 3 shows the brain areas where we found significant differences between neurotypicals and ASD for male (A) and female (B) cohorts separately. For the male cohort, there were seven regions with lower and five regions with larger Ppos coefficients in neurotypical individuals compared to autism. In contrast, within the female cohort, each of the nine areas that showed significant differences had smaller Ppos coefficients in neurotypical individuals compared to those with autism. In the case of Pneg coefficients, both male and female cohorts presented only four areas with significant differences although different ones.

**Fig. 3.**
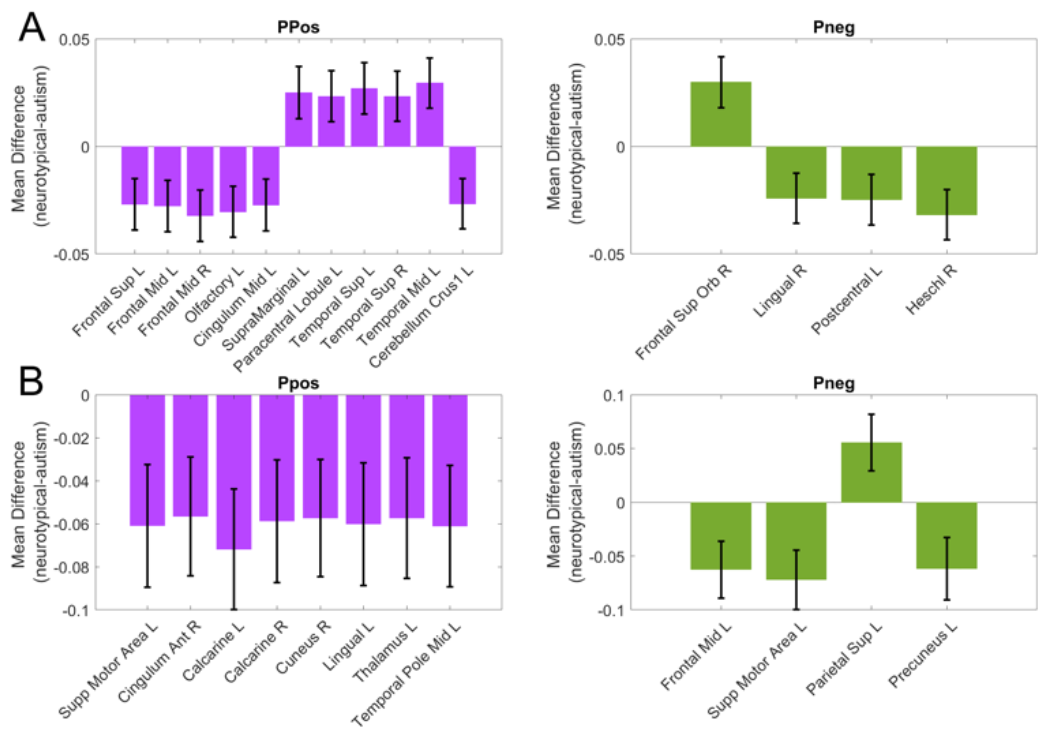
Gender-specific estimation of positive (Ppos) and negative (Pneg) participation coefficients. A) Male Cohort: The bar graphs illustrate the significant mean differences across brain regions for the male group, comparing individuals with autism to neurotypical controls. Error bars indicate the standard error of these estimates, underscoring the variability within each group. B) Female Cohort.

## IV. DISCUSSION

In the pursuit of understanding the complex dynamics of the brain, resting-state fMRI-based brain dynamics models have emerged as a pivotal tool. The model proposed in this study aimed to capture the BOLD signal variations across brain regions, accounting for both linear and nonlinear terms as well as the collective influence of other regions through a term scaled by the parameter *k*_*i*_ and functional connectivity. Despite the advances this model represents, it is not without limitations. For instance, the complexity of neurovascular coupling and the diverse physiological mechanisms that contribute to the BOLD signal [36], [37], [38] are not fully encompassed by our model.

To enhance the BDM of the BOLD-rsfMRI signal, we can consider two significant extensions. First, the current model’s assumption of nonlinearity up to the quadratic order for the area BOLD signal *y*_*i*_ could be expanded to include higher-order nonlinearities. This would allow for capturing more complex and realistic neural responses that extend beyond simple quadratic interactions, which could be pivotal in understanding intricate brain functions and pathologies. Second, the model presently incorporates only linear terms when considering interactions between brain areas. Recognizing that neural interactions are inherently nonlinear and that the connectivity strength can vary with the system’s state [39], one could consider the addition of nonlinear terms to attain a more accurate representation of these dynamics.

Moreover, integrating the modeling of the BOLD signal from rsfMRI data with multimodal neuroimaging data such as Electroencephalography (EEG) and diffusion weighted magnetic resonance imaging (DWMRI) could lead to a more holistic representation of brain dynamics [8], [9]. The combination of these modalities, each contributing unique and complementary information, could facilitate a more accurate depiction of the neural dynamics and network states underlying conditions such as ASD [40]. To realize such a synthesis, advancements in computational methodologies, potentially extending the capabilities of the PINN framework, will be necessary to manage the complex, multimodal datasets [41], [42]. Developing and validating these integrative models against diverse neuroimaging data will ensure that they not only capture the complexity of brain dynamics but also align with empirical observations from multiple perspectives [43].

The modeling of the BOLD signal we present in this study (see (1)) diverges fundamentally from DCM [6] or neural mass models [8]. While DCM is categorized as a state-space model— detailing neuronal activity through differential equations to depict neuronal and synaptic interactions, subsequently linked to BOLD signals via an observational model often comprising a hemodynamic component [44] —our model takes a different stance. It directly models the BOLD signal dynamics, bypassing the explicit distinction between neuronal states and observed BOLD responses. Our differential equations are crafted to define the BOLD signal in relation to its own activity and inter-regional brain interactions, modulated by parameters *a*_*i*_, *b*_*i*_, and *k*_*i*_. By doing so, it aims to capture both the direct and indirect effects that neuronal activity has on the BOLD signal.

The utilization of PINN in this paper offers a novel approach to address current challenges in computational neuroscience and neuroimaging by embedding physical laws into the learning process of neural networks. However, the PINNs framework is not without its limitations [13], [41]. The generalizability of PINNs across the heterogeneous landscape of brain dynamics can be limited, and specific applications may be constrained by computational resources and the chosen network architecture. The assumptions made in this study regarding noise in the fMRI data also present a potential limitation, as during the training we assumed the absence of noise, suggesting that signal variations are solely due to inter-regional interactions. To extend the PINN framework to account for the intrinsic noise in fMRI data, future work could involve the integration of stochastic differential equations within the training process. This would enable the network to learn the variability of the data, not just the central tendencies. Incorporating stochastic elements into PINNs [45] could provide a more comprehensive understanding of brain dynamics, capturing both the deterministic and random aspects of neural signaling.

The findings from this study, while highlighting the potential of PINNs in neuroscience, prompt further research into enhancing the model’s capability to interpret the BOLD signal by incorporating additional physiological measurements and refining the neural network architecture to capture the full complexity of brain dynamics. As we continue to unravel the complex neural signatures of disorders like autism, the convergence of PINNs with advanced neuroimaging analysis is set to play a crucial role in the advancement of computational neuroscience.

## ACKNOWLEDGMENT

The authors are grateful for access to the Tier 2 High-Performance Computing resources provided by the Northern Ireland High Performance Computing (NI-HPC) facility funded by the Engineering and Physical Sciences Research Council (EPSRC), Grant No. EP/T022175/1.

## Footnotes

This work was supported by Grant 222300868 from the Alberta Innovates LevMax program, and by RGPIN-2022-03042 from Natural Sciences and Engineering Research Council of Canada.

